# Changes in searching behaviour of CSL transcription complexes in Notch active conditions

**DOI:** 10.1101/2023.08.10.552734

**Authors:** Sarah Baloul, Charalambos Roussos, Maria Gomez-Lamarca, Leila Muresan, Sarah Bray

## Abstract

During development cells receive a variety of signals, which are of crucial importance to their fate determination. One such source of signal is the Notch signalling pathway, where Notch activity regulates expression of target genes through the core transcription factor CSL. To understand changes in transcription factor behaviour that lead to transcriptional changes in Notch active cells, we have probed CSL behaviours in real time, using *in vivo* Single Molecule Localisation Microscopy. Trajectory analysis reveals that Notch-On conditions increase the fraction of bound CSL molecules, but also the proportion of molecules with exploratory behaviours. These properties are shared by the co-activator Mastermind. Furthermore, both CSL and Mastermind, exhibit characteristics of local exploration near a Notch target locus. A similar behaviour is observed for CSL molecules diffusing in the vicinity of other bound CSL clusters. We suggest therefore that CSL acquires an exploratory behaviour when part of the activation complex, favouring local searching and retention close to its target enhancers. This change explains how CSL can efficiently increases its occupancy at target sites in Notch-ON conditions.

## Introduction

Animal development is shaped by the activity of several highly conserved signalling pathways (Ingham and McMahon 2001; Wiese, Nusse, and van Amerongen 2018; Herrera and Bach 2019). One such pathway is the Notch signalling pathway which, through the core transcription factor CSL (<u>C</u>BF1, <u>S</u>uppressor of Hairless, <u>L</u>ag), activates cohorts of target genes to regulate decisions during the development of many tissues (Artavanis-Tsakonas, Rand, and Lake 1999; Bray 2006; Kopan and Ilagan 2009; Kovall and Blacklow 2010; Bray 2016). As the dysfunction of the pathway has been linked with various human pathological contexts (Ntziachristos et al. 2014; Nowell and Radtke 2017), there are stakes in understanding the molecular mechanisms that enable CSL complexes to find and interact with their target loci to implement Notch pathway activity.

The primary link between cell signalling and gene regulation are transcription factors, the behaviour of which is affected by signal activation through many different mechanisms. Several of these involve regulated nuclear translocation, enabled by a release from cytoplasmic tethers or by association with a co-factor (Weidemüller et al. 2021). In the case of the Notch pathway, activation brings about cleavage of the Notch receptor to release the intracellular domain, NICD, which translocates into the nucleus (Schroeter, Kisslinger, and Kopan 1998; Gordon et al. 2007). There it forms an activator complex with CSL and co-activator Mastermind (Petcherski and Kimble 2000; Nam et al. 2006; Wilson and Kovall 2006). However, in the absence of Notch activity, CSL is present in repressor complexes, partnered with a variety of co-repressors that include co-repressor Hairless in *Drosophila* (Morel et al. 2001; Barolo et al. 2002; Yuan et al. 2016). The activator and repressor complexes co-exist in the nucleus and, although both types of complex have the same DNA binding properties in vitro (Wilson and Kovall 2006; Bianco et al. 2010), signalling leads to enrichment of CSL activation complexes at target enhancers (Krejčí and Bray 2007; Castel et al. 2013; Wang et al. 2014; Gomez-Lamarca et al. 2018; deHaro-Arbona et al. 2023). One question therefore is how the properties of CSL are altered to favour recruitment in Notch active conditions and what mechanisms influence the target search and binding processes.

Knowledge about the fundamental diffusive motion of transcription factors has been proven informative for understanding their recruitment to and interactions with target enhancers in many different pathways and systems (Izeddin et al. 2014; Woringer and Darzacq 2018; Stavreva et al. 2019; Hansen et al. 2020; Tang et al. 2021; Mazzocca et al. 2023). Live-cell imaging has revealed that interactions with cognate binding sites typically last seconds and that transcription factors exhibit a range of dynamic behaviours. For example, both CTCF and CBX2 manifest properties consistent with local confined motion that could reduce the time taken to find their target regulatory elements (Hansen et al. 2020; Kent et al. 2020). In addition, the transcription factors FOXA1 and SOX2 exhibited distinct chromatin-scanning dynamics that related to their searching behaviours (Lerner et al. 2023). It is therefore possible that CSL activation complexes acquire different searching or scanning behaviours which enable them to become efficiently recruited to target enhancers.

To investigate the diffusive properties of CSL in real-time and measure the changes caused by Notch activation we performed Single Molecule Localisation Microscopy, SMLM, (Mazza et al. 2012; Gebhardt et al. 2013) of endogenous CSL, co-activator Mastermind, and co-repressor Hairless in live tissues. Our results show that Notch activation leads to increased exploration and more anisotropic CSL behaviour, consistent with the properties we detect for Mastermind, its partner in the activation complex. The increase in CSL exploratory behaviour is more apparent in proximity to a target locus, *E(spl)-C*, where it becomes locally restricted. Moreover, as the distribution of CSL dynamics is not spatially homogeneous - with bound molecules clustered in different parts of the nucleus and anisotropic behaviour being more common near these clusters - we suggest that in Notch-On conditions, the local exploration that CSL activation complexes undergo near *E(spl)-C* occurs at many different regions in the nucleus, and is a mechanism that enables them to locate their target sites more efficiently.

## Results and Discussion

### Diffusion dynamics of CSL in Notch-Off conditions

We first set out to characterize the properties and analyse the kinetic dynamics of CSL in nuclei without endogenous Notch pathway activity using SMLM. We generated a Halo-tagged CSL (Halo::CSL), expressed at endogenous levels from a genomic rescue construct (Gomez-Lamarca et al. 2018) and tracked its behaviour in *Drosophila* larval salivary glands, whose large nuclei and polytene chromosome make them amenable for live imaging of transcription factors (Lis 2007). To resolve single molecules of CSL, salivary glands were incubated with a limiting concentration of the Halo ligand TMR and imaged with 10ms and 50ms exposure times for 3 to 6 minutes (Supplementary Movie 1). For comparison, we also carried out SMLM of Halo-labelled Histone H2AV in the same tissue (Supplementary Movie 2).

Following SMLM movie acquisition, we performed single molecule localisation using a Gaussian fitting-based approach (Ovesný et al. 2014). Consecutive frame localisations likely to represent the same molecule were then linked into trajectories, by employing a Multiple Hypotheses Tracking Icy-plugin (Chenouard, Bloch, and Olivo-Marin 2013) (Fig 1A). To analyse the resulting trajectories we utilised a variational Bayesian treatment of Hidden Markov models (vbSPT) (Persson et al. 2013) to assign them in one of *n* populations, each defined by a unique Brownian motion diffusion coefficient. We first partitioned trajectories into two populations. With 10ms exposure times, the slow-moving population accounted for 34% of CSL trajectories in comparison to 62% of H2AV trajectories, indicating that a relatively smaller proportion of CSL are stably retained on the chromatin. With the 50ms exposure time, a greater proportion of the fastest moving molecules would not be detected as they become blurred. As expected, the average diffusion coefficient of the fastest molecules population was concomitantly reduced from 1.192µm^2^/s to 0.367 µm^2^/s for CSL (Fig 1B and Supplementary Movie 3), indicating that the longer exposure times will give more scope for teasing apart different properties among the molecules whose motions are slowed by their interactions with the chromatin, making it possible to distinguish exploratory behaviours.

**Figure 1:**
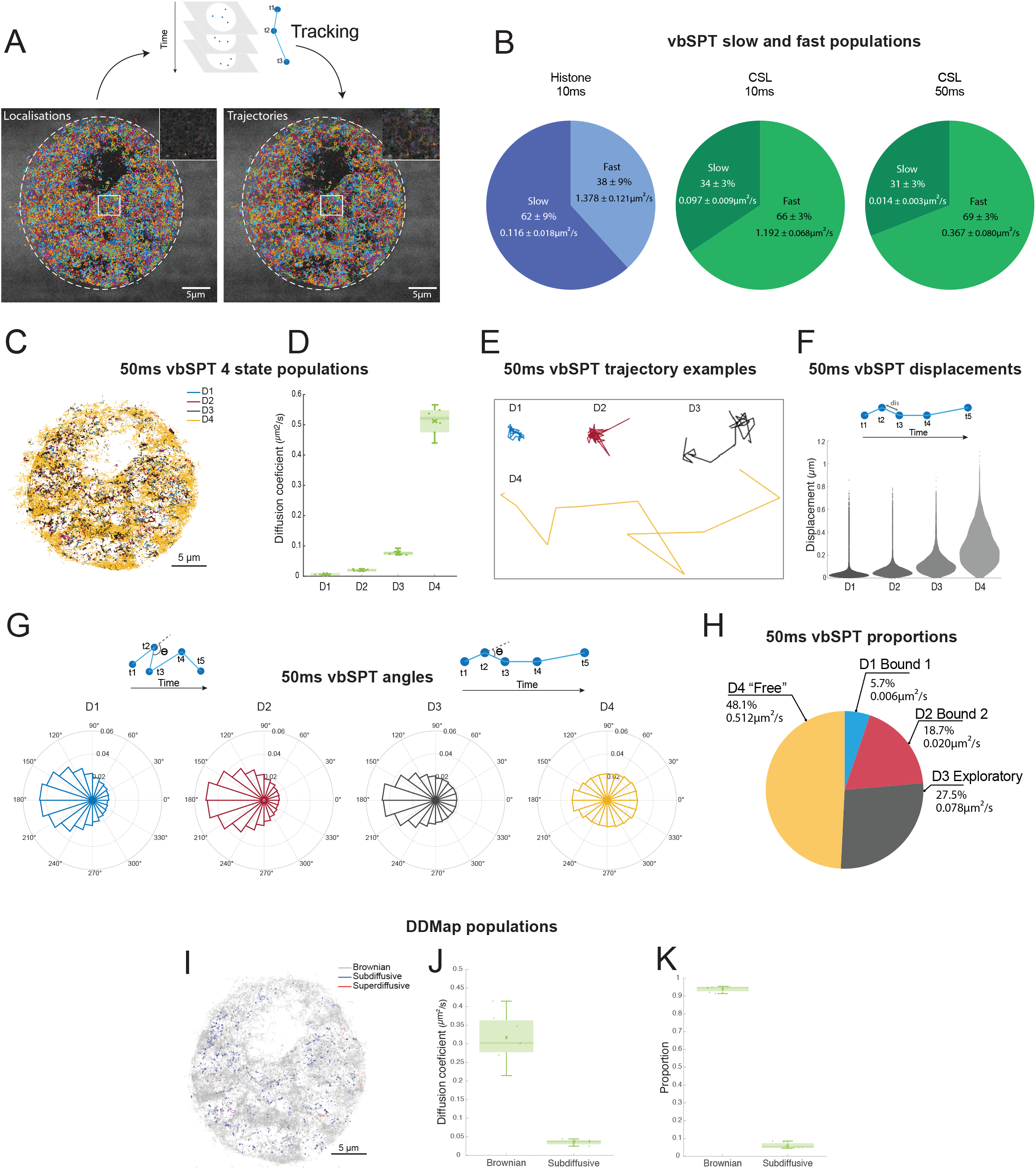
Characterising different types of CSL behaviour in Notch-Off conditions using SMLM analysis pipeline. **A.** Example of single molecule localisations of CSL::Halo acquired at 50ms exposure (left) and of the trajectories defined by tracking the same molecule in consecutive time frames (right), as illustrated in the schematic above. Dotted lines outline nucleus and insets are 10x zoom of the representative region annotated by the white rectangle. **B.** Mean proportion of Histone (H2av) and CSL trajectories belonging to each of 2 populations (slow and fast) defined by vbSPT analysis from 10ms and 50ms exposure times with mean diffusion coefficients. 10ms histone n = 3 nuclei, 22820 trajectories; 10ms CSL, n = 3 nuclei, 35992 trajectories; 50ms CSL n = 7 nuclei, 72980 trajectories. **C.** Nuclear localisations of single molecule CSL trajectories (representative 50ms experiment), coloured according to vbSPT diffusion state: blue D1, red D2, black D3, yellow D4. **D.** Average diffusion coefficients of 4 CSL populations assigned by vbSPT analysis of 50ms movies, n=7 nuclei. See Table S2 for mean values (± SD). For all boxplots, line indicates median, crosses represent mean, boxes 5–95 percentiles and whiskers upper and lower extrema. Data from these 7 nuclei form the basis for analysis in F-K. **E.** Representative examples of trajectories from indicated vbSPT populations. **F.** Distribution of displacements (*dis*) per population, measurements of *dis* the distance between two consecutive localisations in a trajectory as illustrated in the schematic. Trajectories from n = 7 nuclei in D (72980 trajectories), were pooled. Displacement value dis= 0.035 ± 0.040μm for D1, dis= 0.061 ± 0.050μm for D2, dis= 0.123 ± 0.088μm for D3 and dis= 0.271 ± 0.155μm for D4. **G.** Circular histograms show angle distributions per population, angle *θ* is measured between three consecutive localisations in a trajectory as illustrated in the schematics, pooling trajectories as in F. The resultant vector, R, gives a measure of accumulation of angles around the mean value (high er R value shows more accumulation). Mean = 176°, 179°, 179°, 137° and R = 0.300, 0.331, 0.217, 0.008 for D1, D2, D3 and D4 respectively. **H.** Pie chart showing average percentages of CSL molecules in each vbSPT population and diffusion coefficient. For D1 to D4 respectively, mean values (± SD) of proportions are 5.7 ± 0.03%, 18.7 ± 0.03%, 27.5 ± 0.10% and 48.1 ± 0.10%. Mean values (± SD) of diffusion coefficients are 0.006 ± 0.001µm^2^/s, 0.020 ± 0.002µm^2^/s, 0.078 ± 0.008µm^2^/s, and 0.512 ± 0.050µm^2^/s. D1-D4 populations are characterised as Bound1, Bound2, Exploratory and Free respectively, based on diffusion coefficients, displacement, and angle analyses. **I.** Nuclear localisations of CSL single molecule trajectories from C coloured according to motion type assigned by DDMap analysis: Brownian (grey), sub-diffusion (blue) and super-diffusion (red). **J.** Average diffusion coefficients of trajectories obtained from DDMap analysis of 7 nuclei as in D, grouped by motion type (means: 0.317 ± 0.066µm^2^/s for Brownian, 0.036 ± 0.007µm^2^/s for sub-diffusive). **K.** Average proportion of trajectories per nucleus of each motion type obtained from DDMap analysis (means: 0.940 ± 0.016 for Brownian, 0.060 ± 0.016 for sub-diffusive).

In order to probe further into possible exploratory behaviours manifest by CSL in Notch-Off conditions, the trajectories from the 50ms exposure times were partitioned into four diffusion states (D1-D4, slowest to fastest diffusion constant) and mapped back onto the nuclei (Fig 1C-D). Some regions were more strongly enriched for D4 fast-moving molecules and likely correspond to regions with lower density of chromatin (Fig 1C). Others were biased towards slower moving molecules although these were widely distributed in clusters throughout. To further characterise these populations, we analysed displacement (Fig 1F) and angle distributions (Fig 1G). Displacement distributions inform about the range of motion, with small displacements indicating a restricted behaviour, as was the case for the slowest populations, D1 and D2 (Fig 1F). The distribution of angles indicates the extent to which the movement differs from the uniform distribution expected for regular Brownian motion. For example, an accumulation of angles around 180°, known as backwards anisotropy, occurs when molecules are more likely to move backwards than forwards. This compact behaviour suggests constraint or “trapping” of the molecules and was more enriched in the D1 and D2 populations (ben-Avraham and Havlin 2000; Liao et al. 2012; Burov et al. 2013; Izeddin et al. 2014).

By combining all these parameters together (diffusion coefficients, displacement, and angle analyses) we identified behaviours characteristic of each population. Those with the smallest diffusion coefficients, D1 and D2, have properties of bound/trapped molecules, with a high degree of anisotropy and little spatial displacement (Fig 1D-G). They accounted for 6% and 19% of the total CSL molecules respectively (Fig 1H). The slightly longer displacements exhibited by D2 molecules (Fig 1F) likely reflect an association with more mobile chromatin and/or local exchange on and off the DNA. The fastest population, D4, accounting for 48% of trajectories, showed properties of more freely diffusing molecules, with less backwards angular anisotropy and large spatial displacements (Fig 1D-G). As noted above, many freely/fast moving molecules in the nucleus will not have been captured with the 50ms regime. Nevertheless, we refer to the D4 population as “free’ as their movements are not demonstrably constrained (Fig 1D-G). Finally, molecules in the D3 population have intermediate characteristics, with less backwards anisotropy and more displacement than the bound populations (Fig 1D-G). These account for 27% and their characteristics suggest an exploratory behaviour, with a mixture of binding/trapping and freer movements (Fig 1E,H).

For comparison we utilised DDMap, a trajectory analysis software that assigns a different diffusion coefficient to each trajectory (Fig 1J) and, using a nonparametric three-decision statistical test, classifies them as either Brownian, sub-diffusive or super-diffusive (Saxton 1993; Briane, Kervrann, and Vimond 2018; Salomon et al. 2020) (Fig 1I). Only 6% of CSL trajectories were classified as sub-diffusive, a proportion close to the one classified as bound D1 with vbSPT analysis. The majority of CSL molecules (94%) were identified as Brownian, suggesting that all other CSL molecules have some element of Brownian motion associated with their trajectories (Fig 1K). Not unexpectedly, a negligible number of trajectories were classified as super-diffusive.

Together the results demonstrate that a relatively small proportion of CSL is stably associated with chromatin in the absence of Notch activity. The majority of CSL molecules are more transiently associated with chromatin and/or freely moving.

### Notch activity leads to increased binding and exploratory behaviour of CSL

To investigate the effect of Notch activity on CSL we acquired SMLM movies at 50ms exposure in Notch-On conditions, achieved by expressing a constitutively active form of Notch in the salivary gland cells (Supplementary Movie 4). When the trajectories were partitioned into 4 states using vbSPT, the resulting populations had similar diffusions constants to those in Notch-Off conditions (Fig S1A, Table S2A), demonstrating that the overall properties were not substantially altered. However, the proportions mapping to each state differed significantly. First, Notch activation led to an increase in the proportion of bound CSL molecules (D1 increased from 6% to 9%, and D2 to from 19% to 22%, Fig 2A). Second, there was a marked increase in the proportion of CSL molecules exhibiting D3 exploratory behaviours (from 27.5% to 37%). Finally, the proportion of “free” molecules was reduced (from 48 to 32%) as a consequence (Fig 2A, Table S3A).

**Figure 2:**
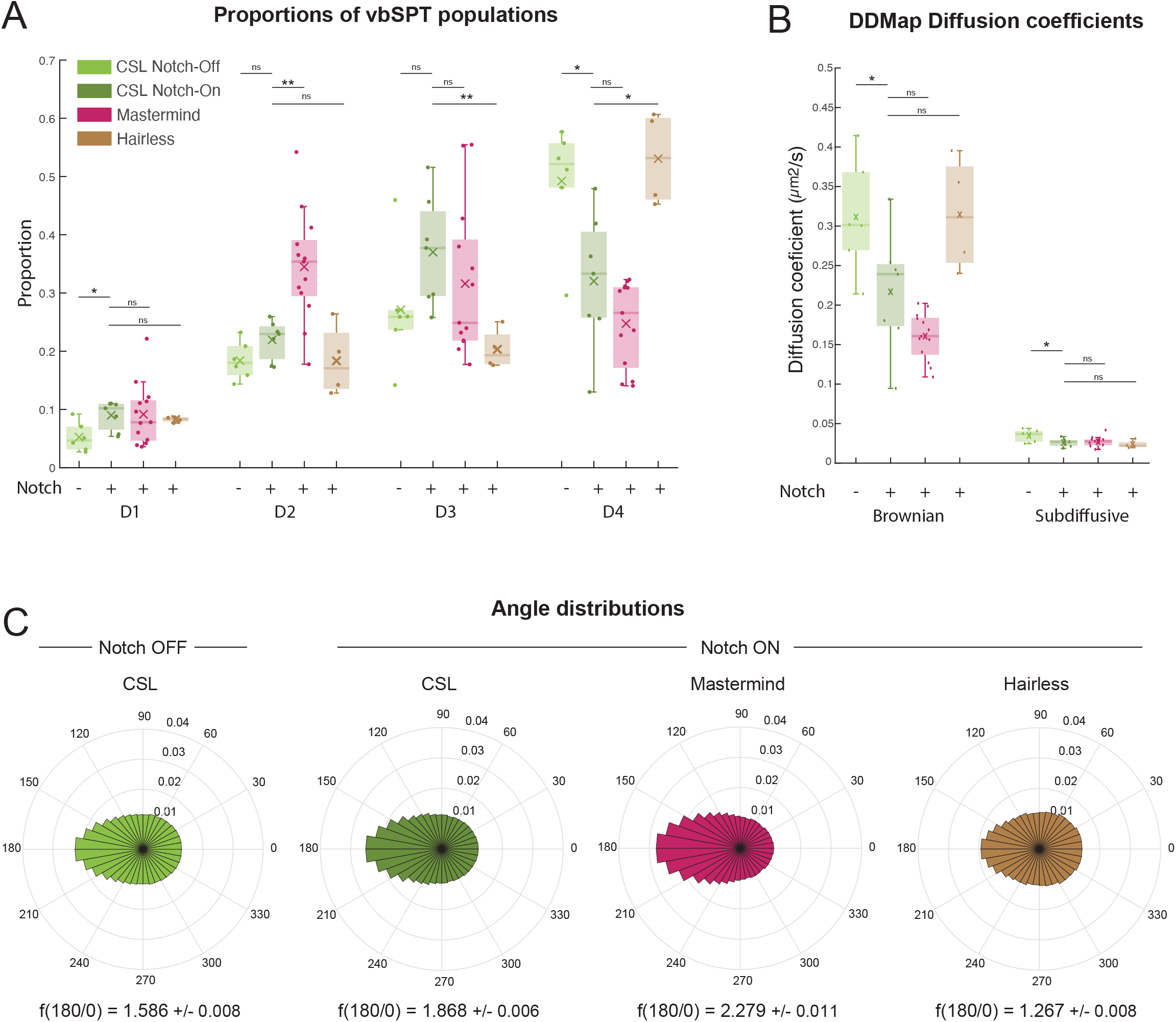
Changes in the dynamics of CSL complexes in Notch active conditions. **A.** Average proportions per nucleus of CSL (green), Mam (red) and Hairless (brown) molecules in each of 4 vbSPT populations in Notch-Off (-) or Notch-On (+) conditions as indicated (CSL Notch-Off n = 7 nuclei, 72980 trajectories; CSL Notch-On n=7 nuclei, 125071 trajectories; Mam n = 13 nuclei, 66753 trajectories; and Hairless n = 4 nuclei, 49657 trajectories). For mean values see Table S1A. Significance was assessed with Mann-Whitney U tests, * p<0.05, ** p<0.01, *** p<0.001. See Table S3A for all p-values. **B.** Average DDMap diffusion coefficients per nucleus in A of Brownian and sub-diffusive CSL (green), Mam (red) and Hairless (brown) molecules in Notch-Off (-) or Notch-On (+) conditions as indicated. For mean values see Table S1B. Significance was assessed with two-sample t-tests, * p<0.05, ** p<0.01, *** p<0.001. For p-values see Table S3B. **C.** Circular histograms of angles, calculated as in Figure 1G, from pooled D3 & D4 CSL trajectories (assigned by vbSPT) in Notch-Off and Notch-On, and from Mastermind and Hairless in Notch-On (nuclei from A). n = 220711, 328143, 142760 and 121396 angles respectively. The fold anisotropy metric f(180/0) ± SD is noted in each case (for more information on f(180/0) see Methods).

The increase in CSL binding was also reflected in the results from DDMap analysis, with the proportions of sub-diffusive trajectories increasing (Fig S1C). More striking were the changes in mean diffusion coefficients which decreased significantly in Notch-On conditions (Fig 2B). These changes are consistent with CSL becoming less freely diffusing and instead exhibiting more stable binding behaviour and more exploration of chromatin in Notch-On conditions, as suggested by the results from the vbSPT analysis.

CSL participates in two types of complexes. In Notch-Off conditions, CSL partners with Hairless to form a corepressor complex (Morel et al. 2001; Barolo et al. 2002; Yuan et al. 2016). In Notch-On conditions, a fraction of CSL molecules are recruited into a tripartite complex with NICD and the co-activator Mastermind (Petcherski and Kimble 2000; Nam et al. 2006; Wilson and Kovall 2006). It is likely that the change in CSL behaviours is related to the nature of the complexes present in the two conditions. We therefore set out to investigate whether the dynamics of Mastermind and Hairless correlated with the different behaviours detected for CSL, by generating endogenously expressed Halo-tagged variants of the two proteins and tracking them with SMLM (Supplementary Movie 5 and 6). The trajectories were partitioned into 4 states using vbSPT, and in all cases the resulting populations had similar diffusions constants consistent with them participating in the same complexes (Fig S1A-B, Table S2A). The characteristics of Hairless were very similar to those of CSL in Notch-Off conditions, (Fig 2A), with a relatively large proportion of D4-freely diffusing molecules (48%). Conversely, behaviour of Mastermind was more closely aligned with that of CSL in Notch-On conditions, with lower proportion of D4 molecules (25%) and a higher proportion of bound molecules (44%). It also exhibited similar proportions with D3, “exploratory”, behaviour (32% Mam vs 37% CSL). Thus, the Notch-induced changes in CSL behaviour are compatible with it acquiring the characteristics associated with its partner Mam, albeit the proportions of Mam that exhibited “bound” D1/D2 behaviours (44%) were even greater than for Notch-On CSL (31%).

To investigate further the change in CSL behaviour in Notch-On conditions, we focused on the more dynamic molecules in the D3 and D4 states and measured their anisotropy to ascertain whether their movement becomes more compact, indicative of a change in searching behaviour (Fig 2C). Indeed, there was an increase in the backwards anisotropy of CSL trajectories in Notch-On conditions, effectively reducing the size of the space it explores. As with the overall dynamics, these properties were more similar to those of Mam than of Hairless - the latter having little backwards anisotropy.

These comparisons show that the behaviour of CSL correlates with those of its partners and shifts from more freely diffusing characteristics in Notch-Off conditions, similar to Hairless behaviour, to more bound and compact behaviours similar to Mastermind in Notch-On conditions.

### Notch-On conditions promote recruitment and exploration at a target locus

The global changes in the properties of CSL could in theory increase the rate at which it binds to target sites. To investigate its movement and recruitment in relation to a specific genomic locus that it regulates, we focussed on the *Enhancer of Split Complex* (*E(spl)-C*) a highly responsive locus containing eleven Notch-regulated genes (Delidakis and Artavanis-Tsakonas 1992; Knust et al. 1992; Jennings et al. 1994; Bailey and Posakony 1995; Lecourtois and Schweisguth 1995; Lai, Bodner, and Posakony 2000). We have previously generated strains for live imaging *E(spl)-C*, introducing INT sequences into the 3’ end of the complex and combining this with fluorescently tagged ParB which recognizes INT, enabling us to visualise the locus as a clear “band” in salivary gland nuclei (Fig 3A) (Gomez-Lamarca et al. 2018).

**Figure 3:**
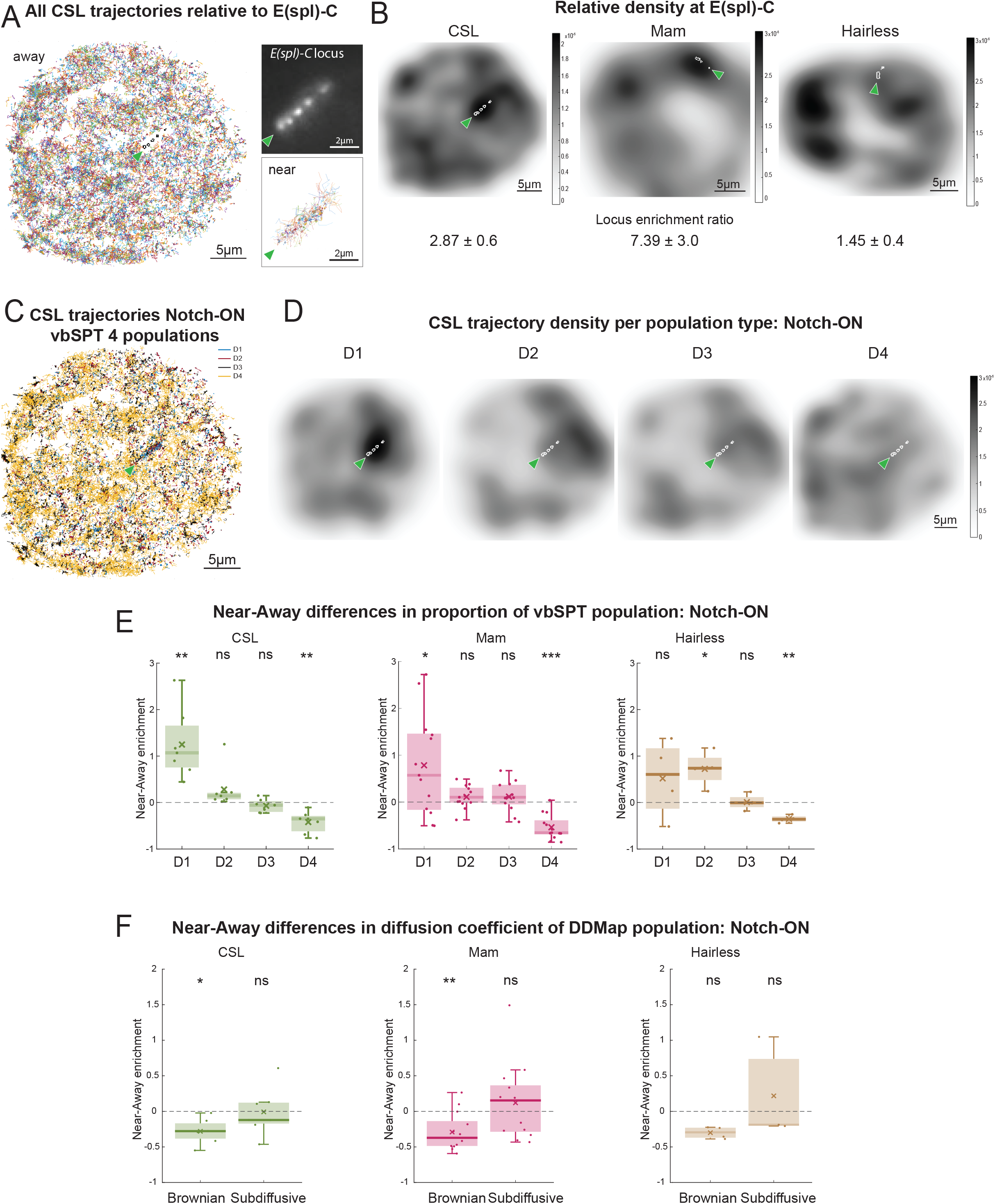
Recruitment and change in behaviour of CSL complexes near *E(spl)-C* in Notch active conditions. **A.** Segregating trajectories relative to *E(spl)-C* (detected by Int/ParB-GFP, arrowhead and upper inset) in a representative Notch-On nucleus. Trajectories localised within a 550nm distance area around *E(spl)-C* locus were classified as ‘near’ (lower inset). Remaining trajectories in the nucleus were classified as ‘away’ (main image). Arrowhead indicates location of *E(spl)-C* locus. **B.** Density map of CSL, Mam and Hairless trajectories from representative Notch-On nuclei. Colourbar is in units of number of trajectories per nm². Arrowhead indicates *E(spl)-C* locus. Mean locus enrichment ratio (± SD) is shown (see Methods). A ratio >1 indicates molecule density near *E(spl)-C* is higher than overall nuclear density. CSL n=7 nuclei, 125071 trajectories, Mam n=13 nuclei, 66753 trajectories and Hairless brown, n=4 nuclei, 49657 trajectories. **C.** Single molecule trajectories from a representative CSL SMLM experiment in a Notch-On nucleus, plotted at their respective localisation positions, colour coded according to vbSPT diffusion state (Blue D1, Red D2, Black D3, Yellow D4). Green arrowhead indicates location of *E(spl)-C* locus. **D.** Density map of each CSL population (D1-D4) from (C) relative to *E(spl)-C* locus (arrowhead). Colour-bar is in units of number of trajectories per nm². **E.** Enrichment of vbSPT populations near *E(spl)-C* for CSL, Mam and Hairless in Notch-On conditions (nuclei as in B). Values plotted were calculated as (Proportion near-Proportion away)/Proportion away, hence a value greater than 0 represents an enrichment of the respective population near the target locus. Significance was assessed using two-sample t-tests, * p<0.05, ** p<0.01, *** p<0.001 and see Table S3D for all p-values. **F.** Relative diffusion coefficients of Brownian and sub-diffusive trajectories near *E(spl)-C* versus away for CSL in Notch-On conditions (nuclei as in B). Values plotted were calculated as (Diffusion coefficient near-Diffusion coefficient away)/Diffusion coefficient away, hence a value smaller than 0 indicates a decrease in the diffusion coefficient near the locus. Significance was assessed using Wilcoxon signed rank-sum tests, * for p<0.05, ** for p<0.01, *** for p<0.001 and see Table S3E for all p-values.

Taking a snap-shot of the *E(spl)-C* at the start and end of each movie, we compared the behaviour of CSL around this locus against behaviour elsewhere in the nucleus. From the density maps of CSL trajectories, it was already evident that there was a significant enrichment of trajectories around *E(spl)-C* (Fig 3B). To quantify the locus associated trajectories relative those elsewhere in the nucleus, we set a distance threshold of 550nm around *E(spl)-C* and every trajectory falling within that region was designated ‘near’ while all other trajectories were designated as ‘away’ (Fig 3A). Firstly, we measured the molecule density (number of trajectories per unit area) near the locus and calculated its ratio to the nuclear density. As suggested by the trajectory maps, (Fig 3B) the ratio was significantly higher in Notch-On conditions, confirming that Notch activity promoted recruitment of CSL molecules to the target locus. As expected, Mam also became enriched near *E(spl)-C* in Notch-On condition, indeed to an even greater extent than CSL (Fig 3B). Conversely, no enrichment was detected for the co-repressor Hairless for which there was a relatively small number of trajectories present near the locus (Fig 3B).

To investigate which type of CSL behaviour was affected, we calculated the enrichment for each diffusive population near the locus in Notch-On conditions. Of the 4 populations only the CSL D1 “bound” population showed a robust enrichment, a pattern that was replicated by the Mam populations. This was accompanied by a relative decrease in the proportion of faster moving D4 molecules (Fig 3D-E). Thus, the proportion of bound CSL and Mam was significantly greater at *E(spl)-C* (Fig 3E, Fig S2B). Despite not becoming enriched at the locus (Fig 3B), Hairless also exhibited proportionately more bound-type behaviour, evidenced by increased proportions of D1, D2 molecules and reduced proportion of D4. The increase in binding near the locus for all three molecules was also captured by DDMap analysis, both through a slight enrichment of the sub-diffusive population and through a significant decrease in the diffusion coefficients of Brownian trajectories near this region (Fig 3F, Fig S2C-D).

The increase in bound CSL molecules at *E(spl)-C* in Notch-On conditions is consistent with previous imaging and ChIP data (Krejčí and Bray 2007; Castel et al. 2013; Wang et al. 2014; Gomez-Lamarca et al. 2018). The fact that there is proportionately more binding for all 3 molecules, including the corepressor Hairless, suggests that there is a change in the chromatin environment that facilitates the binding or trapping of the molecules in that region (Gomez-Lamarca et al. 2018; deHaro-Arbona et al. 2023).

### Notch activity promotes local searching by CSL complexes

The increase in binding of CSL at *E(spl)-C* in Notch-On conditions led us to question whether there was a change in its local behaviour to favour its recruitment in the vicinity. For example, an increase in the anisotropy of trajectories, would reflect more compact diffusion around the locus that could aid recruitment. We therefore analysed the anisotropy of D3 and D4 trajectories near *E(spl)-C* in comparison to elsewhere in the nucleus. Based on the anisotropy metric f(180/0), there was a large increase in CSL backwards anisotropy near *E(spl)-C* (2.75 vs 1.85, Fig 4A). This implies that the effective size of the space CSL explores is reduced, by local trapping or transient binding, which would favour its on-rate at the target locus (Slutsky and Mirny 2004; Kapanidis, Uphoff, and Stracy 2018; Woringer and Darzacq 2018; Hansen et al. 2020; Darzacq and Tjian 2022).

**Figure 4:**
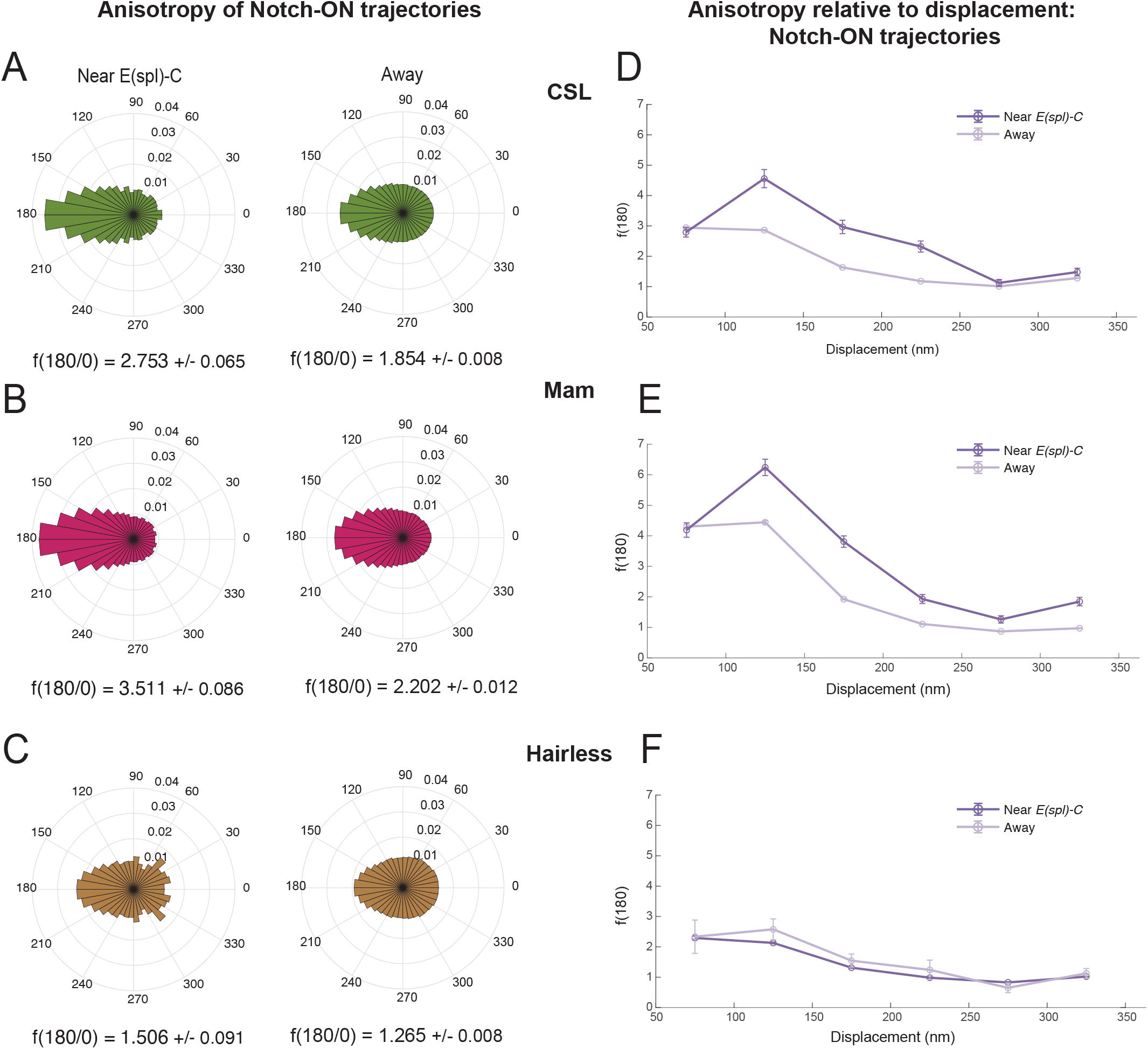
Increased Searching behaviour of CSL complexes at target locus in Notch-On conditions. **A-C.** Circular histograms of angles calculated from pooled D3 & D4 trajectories (assigned by vbSPT) for CSL (A), Mam (B) and Hairless (C) in Notch-On nuclei near *E(spl)-C* locus and away from it (nuclei as in Figure 3A). CSL n = 6158 angles near, n = 321985 angles away; Mam n = 10371 angles near, n = 132389 angles away; and Hairless n = 945 angles near, n = 120451 angles away. The value of f(180/0) ± SD is also given for each distribution. **D-F.** Anisotropy [f(180/0)] relative to molecule displacement (nm) for D3 and D4 trajectories of CSL (A), Mam (B) and Hairless (C) near *E(spl)-C* locus and away in Notch-On conditions (nuclei as in Figure 3A). Error bars show the standard deviation from bootstrapping with 50 iterations (for more information see Methods).

If this model is correct, the anisotropy should vary spatially with respect to the target locus. We therefore asked how anisotropy changed with displacement length, keeping our analysis within a displacement range of 50 – 350nm. This provided accuracy by excluding displacements close to/lower than our localisation error (20nm) (see Methods) and by ensuring sufficient numbers of angles for the longest displacements in the range. First it was evident that the pattern of CSL anisotropy differed near the locus compared to away. Notably, near the locus CSL, acquired a peak of anisotropy between 100-150nm, suggesting that there is a tendency for CSL molecules to become trapped in regions of that size (Fig 4D). Combined with the fact that anisotropy dropped for longer displacements, it implies a form of local exploration near the locus (Slutsky and Mirny 2004; Kapanidis, Uphoff, and Stracy 2018; Woringer and Darzacq 2018; Hansen et al. 2020; Darzacq and Tjian 2022). A similar behaviour was observed for the co-activator Mam, whose anisotropy also displayed a peak at the 100-150nm range (Fig 4B, E). In contrast, Hairless showed no similar peak in anisotropy near the locus for any specific displacements (Fig 4C, F) arguing that the corepressor complex lacks the features that favour local searching.

Overall, our results show that Notch activity brings about changes in the behaviour of CSL at target loci. The properties it acquires, of increased binding and a more restricted compact diffusion, resemble those of Mam, consistent with the two forming a complex. Their anisotropic diffusion close to the target locus implies that they engage in local exploration (Izeddin et al. 2014; Hansen et al. 2020; Mazzocca et al. 2021) that could aid their recruitment.

### Clustering analysis reveals higher backwards anisotropy near clusters of bound CSL

To investigate whether the searching behaviour observed at *E(spl)-C* is a more general feature of Notch-On conditions, we asked whether similar anisotropy was present at other genomic locations where CSL was bound. First, we identified clusters of bound CSL (D1 and D2) molecules in each of our datasets (excluding those at *E(spl)-C*) and then analysed the anisotropy of the more motile populations in relation to those clusters.

Clustering analysis was carried out using a superposition of Poisson process models (Byers and Raftery 1998, see Methods). Using bound trajectory centres as datapoints, the distance of each point to its k^th^ nearest neighbour was calculated. The E-M algorithm was then employed for fitting a mixture distribution to these distances, using a high density (to identify clustered molecules) and a low density (to identify non-clustered molecules) Poisson process. Our analysis remained robust over a range of k-values, but we focussed on k = 11, which identified larger and denser clusters, making these regions plausible candidates for other target enhancers (Fig 5, Fig S3A).

**Figure 5:**
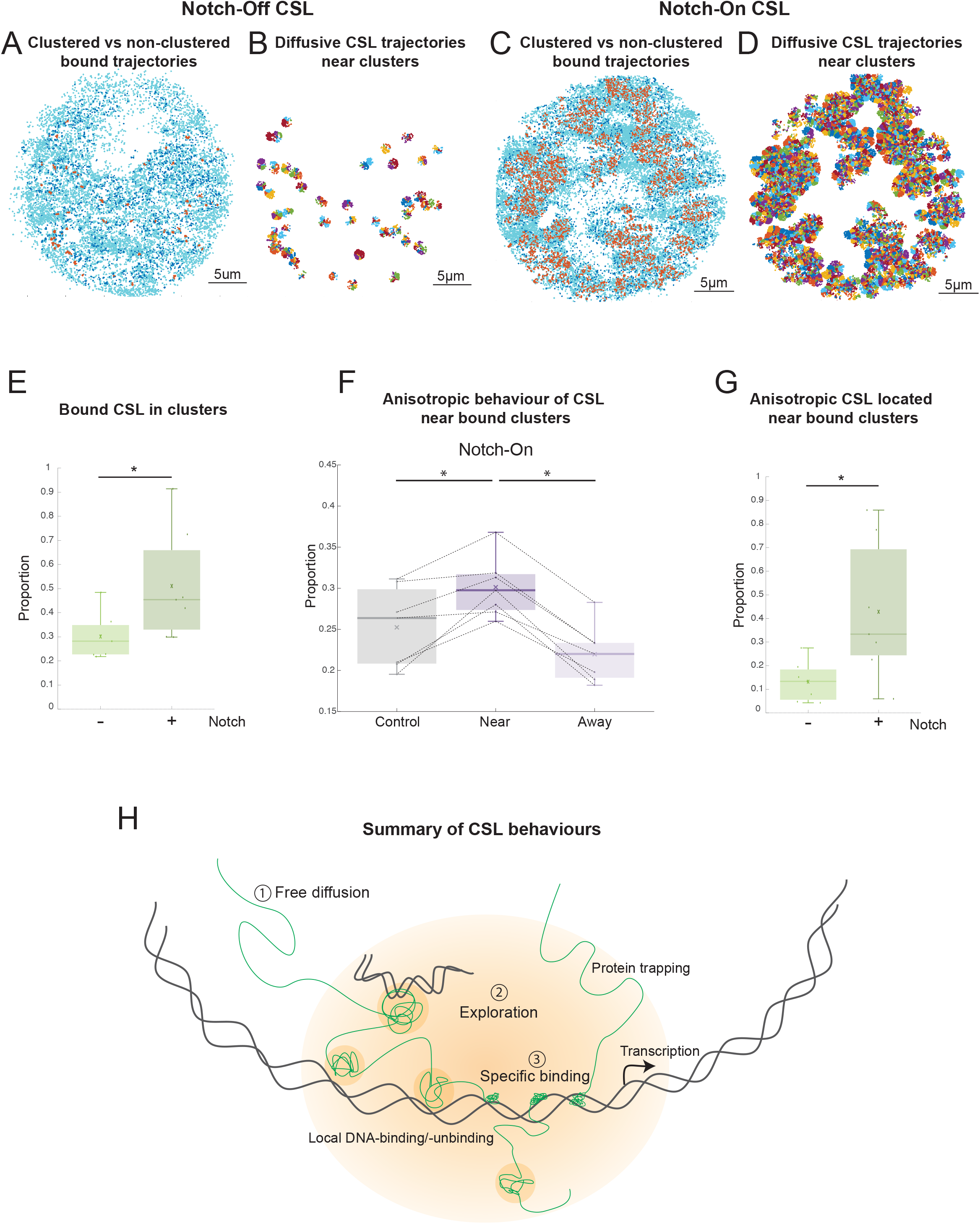
Clustering analysis reveals higher backwards anisotropy near clusters of bound CSL. **A,C**. Clustered bound trajectories identified using a superposition of Poisson process models from representative examples of CSL Notch-Off (A) and Notch-On (C) nuclei. Each dot represents the centre of a trajectory. Orange, clustered bound trajectories; dark blue, bound trajectories identified as non-clustered; light blue, diffusive trajectories (D3 & D4). **B,D.** CSL diffusive jumps (D3 & D4) in nuclei shown in (A) and (C), identified as ‘near-cluster’ using a 550nm distance threshold around each clustered bound trajectory. **E.** Proportion of bound trajectories that were identified as clustered for CSL Notch-Off (0.301 ± 0.096) and Notch-On (0.511 ± 0.228) nuclei. A two-sample t-test was performed, p = 0.045. **F.** Proportion of near and away diffusive jumps (D3 & D4) that were anisotropic in CSL Notch-On nuclei. Mean values: 0.252 ± 0.049 Control, 0.301 ± 0.037 Near, 0.220 ± 0.035 Away. Lines connect data from the same experiment (see Results & Methods for details on control analysis). Wilcoxon signed-rank tests were performed, p = 0.016 for both Control vs Near and Near vs Away. **G.** Proportion of all anisotropic diffusive jumps (D3 & D4) that were identified as ‘near-cluster’ for CSL Notch-Off (0.132 ± 0.084) and Notch-On (0.428 ± 0.291). A two-sample t-test was performed, p = 0.036. **H.** Schematic illustrating trajectories of CSL complexes (green). Away from chromatin, CSL complexes are freely diffusing (1). In Notch-On conditions CSL acquires exploratory behaviour (2) near target enhancer comprised of local binding-unbinding to DNA and reflection from boundaries of high protein concentration (orange). Together these favour increased specific binding (3) at regulated genes.

Clustering of CSL bound molecules was significantly more common in Notch-On nuclei, (Fig 5A, C and E), the proportion of clustered bound molecules was 1.7-fold higher in Notch-On conditions than in Notch-Off. To analyse the behaviour of molecules close to the clusters we set a 550nm search radius and captured all D3 and D4 localisations falling within that region (Fig 5B, D). These constituted what we defined as ‘near-cluster’ jumps, while all other jumps were ‘away-cluster’, and we calculated what proportion of ‘near’ jumps were anisotropic (see Methods). As a control we randomly shifted the position of the clusters and repeated the anisotropy analysis for localisations near the ‘fake’ clusters (see Methods). Our results showed a significantly higher anisotropy in the diffusion properties of molecules near the bone fide CSL clusters compared to those located away from clusters in Notch-On conditions (Fig 5F). Unexpectedly, this difference was also detected in Notch-Off conditions, albeit there were many fewer CSL clusters (Fig S3B). These results suggest that the change in anisotropy is linked to the clustering of bound complexes rather than to a specific property of the activation complexes themselves. One likely explanation is that the bound clusters are associated with regions of more accessible chromatin, which would provide an environment that favoured searching, giving rise to more anisotropic behaviours. While increased anisotropy was observed near bound clusters regardless of Notch activity status, when comparing the proportion of anisotropic displacements that were located near bound clusters, we found this to be much higher in Notch-On conditions (42.8% versus 13.2%) (Fig 5G). If our bound cluster/open chromatin correlation hypothesis is correct, the accumulation of anisotropic behaviour near these clusters in Notch-On could indicate a mechanism through which local searching in regions of open chromatin enables these activation complexes to efficiently find their target sites.

## Conclusion

In summary, tracking the behaviour of CSL molecules in real time reveals that they acquire more exploratory behaviours in Notch active conditions. Characterised by restricted and anisotropic diffusion these behaviours are most evident close to a target gene locus and could aid recruitment and binding at regulated enhancers (Fig 5H). Anisotropy is thought to originate from a combination of two mechanisms. One probable cause is reattachment to the DNA. When a protein dissociates from a site where it is bound, it is more likely to re-attach to the same region rather than binding elsewhere. In the time-scale of SPT experiments (milliseconds) the protein will likely step back following a forward jump. The second mechanism involves protein trapping. If a protein engages in interactions with other factors, it will be reflected back from the boundary of protein rich domains, leading to an increased probability of backward motion. The range of marked anisotropy is indicative of the size of the zone where the protein is trapped (Hansen et al. 2020). The properties are thus consistent with Notch activity promoting formation of a transcription hub that involves a combination of altered chromatin accessibility, increasing the probability of DNA-protein interactions, and multivalent protein interactions, retaining the proteins in proximity (Fig 5H). Similar exploratory behaviour has been detected for factors such as CTCF and CBX2, and has been linked to the formation of clusters or trapping zones, that in some cases equate to condensates (Hansen et al. 2020; Kent et al. 2020). Our results align with these studies and suggest a general mechanism where changes in protein composition and chromatin accessibility, instigated by signalling, promote local searching to enable efficient recruitment of transcription complexes to their target sites.

One limitation of our study is that the *Drosophila* salivary gland is unusual, having multiple genomic copies in its polytene chromosomes. This organisation facilitates the imaging and monitoring of changes with respect to a target gene but may have some unusual properties. For example, there may be more extensive local hopping between the co-aligned DNA strands that could favour more exploratory behaviour. We note also that while we are following endogenous levels of the nuclear factors, Notch activity is provided ectopically. However, our observations are consistent with the increased CSL recruitment detected by ChIP in a range of cell types (Krejčí and Bray 2007; Castel et al. 2013; Wang et al. 2014), making it likely that they reflect general properties of these transcription complexes.

## Methods

### Experimental Animals

*Drosophila melanogaster* flies, as indicated, were maintained at room temperature (approximately 20°C) using standard cornmeal food (Glucose 76g/l, Cornmeal flour 69g/l, Methylparaben 2.5ml/l, Agar 4.5g/l and Yeast 15g/l.) Experimental crosses, from which larvae were collected for dissections, were kept at 25°C.

### Fly stocks and crosses

Notch activity was provided by combining *UAS-NΔECD* (Fortini et al. 1993; Rebay, Fehon, and Artavanis-Tsakonas 1993) with *1151-Gal4* (FBti0007229, Gomez-Lamarca et al. 2018), the control *UAS-LacZ* (FBti0018252) was used in Notch-Off glands. The indicated Halo-tagged lines, as described below, were present in each condition and the *E(spl)-C* locus was labelled with an INT insertion detected by ParB1-GFP (Gomez-Lamarca et al. 2018). Full genotypes are provided in Table S4.

### Generating Halo-tagged protein lines

To generate lines expressing endogenous levels of Su(H)::Halo and Hairless::Halo, we modified the AttB plasmids containing genomic rescue constructs Su(H)-eGFP and Hairless:eGFP, (Gomez-Lamarca et al. 2018). Briefly, eGFP coding sequence was replaced by HaloTag (pFN23K-Halo plasmid, given by St Jonhston lab, Promega G2861), amplified using primers CACCTAGGATGGCAGAAATCGGTACTGGCTTTCCATTCGACC and CTACGCGTTGCCGGAAATCTCGAGCGTGG for Su(H)::Halo, and primers TTACAGATCTCTGAAATCGGTACTGGCTTTCC and TTTCTAGAGACAGCCGGAAATCTCGAGCGTGG for Hairless::Halo. Plasmids and HaloTag PCR products were digested using AvrII (New England Biolabs, R0174S) and MluI (NEB, R0198S) for Su(H)::Halo, or BglII (NEB, R0144L) and XbaI (NEB, R0145S) for Hairless::Halo. The resulting AttB plasmids were injected into a strain containing phiC31 integrase and AttP site in position 86F8 in chromosome 3 (Bloomington 24749, Su(H)) or position 51D in chromosome 2 (Bloomington stock 24483, Hairless) to generate transgenic Su(H)::Halo or Hairless::Halo flies.

Mam::Halo flies were generated using CRISPR/Cas9 genome engineering (flycrispr.org) to insert the coding sequence of HaloTag into the endogenous Mam gene. Briefly, a plasmid containing homology arms flanking the starting codon of Mam, HaloTag, SV40 and the PiggyBac 3xPax3-dsRED cassette from pHD-ScarlessDsRed (flycrispr.org) was synthesized by NBS Biologicals. Transformants were obtained by co-injecting this plasmid with a pCFD3-dU6:3gRNA plasmid (Addgene #49410) expressing the gRNA GACGCATTTATGGATGCGGG and screening for 3xPax3dsRED. The 3xPax3-dsRED cassette was excised by crossing with αTub84B-PiggyBac flies (BDSC #32070). Maps of the homology and gRNA plasmids and final genomic sequence can be found at https://benchling.com/bray_lab/f_/bRaeXvzx-mamhalo-endogenous-tag/.

### Salivary Gland Cultures and Halo ligand Treatments

Salivary glands were isolated from third instar larvae grown at 25°. Dissections and mounting in observation chambers were performed as described (Gomez-Lamarca et al. 2018) except for the additional following steps to sparse label with Halo ligand. Glands were incubated for 15min in TMR Halo ligand (Promega, G825A) diluted in Dissecting medium (Gomez-Lamarca et al. 2018). Concentration of ligand was first adjusted in each case by serial dilutions to reach single molecule resolution which corresponded to a concentration of 10nM for CSL-Halo and Hairless-Halo, of 50nM for Mam-Halo and of 0.01-0.02nM for H2AV-Halo. After incubation the glands were washed 3 × 10min in Dissecting medium and briefly rinsed in PBS(1X) (Thermofisher, #70011044) before mounting.

### Single Molecule Localisation Microscopy

The custom build microscope and localisation errors were as described in Gomez-Lamarca et al. 2018. Samples were continuously illuminated during imaging with a 561nm excitation wavelength laser to excite the Halo ligand emitting at 585nm and a 488nm excitation wavelength laser to excite the GFP-labelled Locus Tag emitting at 510nm. Laser power used for imaging with these lasers was approximately 500W/cm2 and 30W/cm2 respectively. For each data set, the region imaged was focused around the nucleus and consisted of a square of approximately 40µmX40µm dimensions. The pixel size of acquired movies was 110nm. Between 4 to 13 nuclei were imaged for each condition each with 50ms exposure time for 3.3 to 6.7 minutes. For the experiments with 10ms exposure time, 3 nuclei were imaged for 2.8 to 3.3 minutes.

### Trajectory analysis basic pipeline

Following movie acquisition, single molecules were localised with a Gaussian fitting-based approach (Ovesný et al. 2014) which allowed the enhancement of localisation precision to subpixel resolutions of ∼20nm (Gomez-Lamarca et al. 2018). A multiple hypothesis tracking algorithm was employed for tracking of the molecules, allowing no detection gaps within tracks, using an Icy-plugin based on Chenouard, Bloch, and Olivo-Marin 2013. Trajectories consisting of at least 4 time points were then analysed using two different methods. The first method was based on a variational Bayesian treatment of Hidden Markov models (vbSPT) (Persson et al. 2013) and assigned each jump (movement of a molecule between two consecutive time points) within a trajectory to one of n populations, each defined by a unique Brownian motion diffusion coefficient. For this study n=2 and 4 were used. To account for the bias towards slow jumps due to defocalisation (fast molecules leaving the focal plane much quicker than slow ones, Kues and Kubitscheck 2002; Mazza et al. 2012; Hansen et al. 2018), any analysis involving population proportions was carried out using the whole trajectories, assigning to each trajectory the mode state of its jumps.

The second analysis method was based on a Dense Mapping approach (DDMap) (Salomon et al. 2020) and considered a wider class of motion types: Brownian, sub-diffusion and directed motion (Saxton 1993). Each trajectory was classified as one of these motion types using a nonparametric three-decision statistical test and a unique diffusion coefficient was calculated for it. DDMap scripts were adapted so that non-spatially averaged diffusion coefficients were calculated. For both trajectory analysis methods, the target locus in each nucleus was detected using image thresholding with Otsu’s method and a 2-D Gaussian smoothing kernel was used for filtering.

### Angle & anisotropy analyses

Standard MATLAB procedures were used for angle calculation, as shown in Fig 1G. Angular statistics, including the calculation of the resultant vector, R, were carried out using the MATLAB package CircStat (Berens 2009). For anisotropy analyses, the fold anisotropy metric f(180/0) was defined and calculated as how many-fold more likely a molecule is to make a step backwards compared to a step forward,

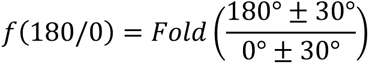

This definition, and scripts required for this analysis were adapted from Hansen et al. 2020. Using only D3 and D4 jumps (as identified by vbSPT analysis) for anisotropy analyses guaranteed the displacements considered to be well above our localisation error (Gomez-Lamarca et al. 2018), ensuring the accuracy of angle calculations. Calculations of f(180/0) against mean displacement were carried out for 50 subsamples using 50% of the data (with replacement). Error bars on Fig 4D-F show SD from these sub-samplings.

### Molecule density analysis

Molecule density maps showing the spatial distribution of molecule trajectories were generated using kernel smoothing in MATLAB, with bandwidth optimised for estimating normal densities (Bowman et al. 1997). The units for all density maps are number of trajectories per 0.0001μm^2^. The design of the maps was carried out using the MATALB package ColorBrewer (DrosteEffect [2015]). Values shown in Fig 3B as “Locus enrichment ratio” represent a fold increase of near-locus density compared to nuclear density and were calculated as follows:

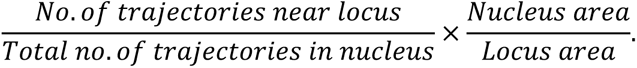

Locus and nucleus areas were calculated with standard MATLAB procedures, using the convex hull of localisations and masking.

### Clustering

Clustering of bound molecules was performed in R using *nnclean*, based on a superposition of Poisson processes model developed by Byers and Raftery, 1998 (Byers and Raftery 1998) and uses the distance of each point to its k^th^ nearest neighbour. For more details on this analysis and the choice of k see Results section. For anisotropy proportions shown in Fig 5F, 5G and Fig S5B, a jump of a D3 or D4 trajectory was regarded as anisotropic if the angle it formed with its successive jump was within the range 180° ± 30°.

For each experiment, control analysis was performed in MATLAB by first identifying a region of interest using the convex hull of diffusive (D3 and D4) jumps. The position of the bound molecules was then randomly shifted in the nucleus (all bound clusters translated by the same vector). If more than 80% of these molecules remained inside the region of interest after translation, anisotropy analysis as described in Results was carried out for the diffusive molecules near & away the shifted clusters. The process was repeated 100 times for each movie and mean values were calculated, which are shown in Fig 5F and Fig S5B.

### Boxplots and Statistical tests

For all boxplots, lines across represent the median, crosses represent the mean, boxes 5–95 percentiles and whiskers upper and lower extrema.

For statistical tests involving only one sample, one sample t-tests (for normal samples) and Wilcoxon rank-sum tests (for not normal samples) were performed. For statistical tests involving two samples, two sample t-tests (if samples were normal) and Mann-Whitney U tests (if samples were not normal) were performed. For paired data (clustering analysis in Fig 5), paired t-tests (if samples were normal) and Wilcoxon signed rank tests (if samples were not normal) were performed. Normality of the samples was assessed using Q-Q plots and Shapiro-Wilk tests. Where two samples were compared, equality of variance was also assessed with Bartlett’s test (if samples were normal) and Levene’s test (if samples were not normal). In all cases significance was presented as follows: * for p<0.05, ** for p<0.01, *** for p<0.001. All statistical tests were performed in R.

## Supporting information

Movie 4 CSL Notch ON 50ms

## Acknowledgments

We thank Kevin O’Holleran, Martin Lenz and Cambridge Advanced Imaging Centre for their advice and help with the imaging, Kat Millen for embryo injections to generate the Halo-tagged proteins, all members of the Bray lab for helpful discussions. The work was funded by a Wellcome Trust Investigator Award (212207/Z/18) to SJB and by an ESPRC Fellowship to LM. CR and SB were supported by studentships from Wolfson College-Dept of Physiology Development and Neuroscience-School of Biological Sciences (University of Cambridge).

## Author contributions

SB, CR, MGL, LM and SJB designed the experiments and analysis. SB acquired the data. MGL generated critical reagents. SB, CR, SB, LM and SJB analysed the data. SB, CR and SJB wrote the manuscript, MGL and LM reviewed and commented on the manuscript.

