## Supplementary Material for "Changes in searching behaviour of CSL transcription complexes in Notch active conditions"

**Supplementary Figures:**

Supplementary Figure S1: Effect of Notch activation on vbSPT diffusion coefficients and DDMap proportions of CSL complexes.

Supplementary Figure S2: Recruitment and change in DDMap proportions of CSL complexes near *E(spl)-C* in Notch active conditions.

Supplementary Figure S3: Clustering analysis – nearest neighbour histogram and anisotropic behaviour of CSL near bound clusters in Notch-Off conditions.

**Supplementary Tables:**

Table S1: Proportions of molecules assigned to each population

Table S2: Diffusion constants.

Table S3: Results from statistical tests.

Table *S*4: Genotypes of flies used for each condition

**Supplementary Movies:**

**Supplementary Movie 1: CSL Notch-Off 10ms**
CSL-Halo sparsely labelled with TMR ligand (10nM) and imaged at 10ms exposure time in a salivary gland nucleus where Notch signalling is inactive.

**Supplementary Movie 2: Histone Notch-Off 10ms**
H2AV-Halo sparsely labelled with TMR ligand (0.01nM) and imaged at 10ms exposure time in a salivary gland nucleus where Notch signalling is inactive.

**Supplementary Movie 3: CSL Notch-Off 50ms**
CSL-Halo sparsely labelled with TMR ligand (10nM) and imaged at 50ms exposure time in a salivary gland nucleus where Notch signalling is inactive.

**Supplementary Movie 4: CSL Notch-On 50ms**
CSL-Halo sparsely labelled with TMR ligand (10nM) and imaged at 50ms exposure time in a salivary gland nucleus where Notch signalling is ectopically activated.

**Supplementary Movie 5: Mastermind Notch-On 50ms**
Mam-Halo sparsely labelled with TMR ligand (50nM) and imaged at 50ms exposure time in a salivary gland nucleus where Notch signalling is ectopically activated.

**Supplementary Movie 6: Hairless Notch-On 50ms**
Hairless-Halo sparsely labelled with TMR ligand (10nM) and imaged at 50ms exposure time in a salivary gland nucleus where Notch signalling is ectopically activated.

#### Supplementary Figure Legends

**Supplementary Figure S1: Effect of Notch activation on vbSPT diffusion coefficients and DDMap proportions of CSL complexes.**

**A.** Diffusion coefficients per nucleus of each vbSPT population for CSL, Mam and Hairless in Notch-Off conditions. See Table S2A for mean values. No significant differences between the different molecules for all four states, using Mann-Whitney U tests.

**B**. Diffusion coefficients per nucleus of each vbSPT population for CSL, Mam and Hairless in Notch-On conditions. See Table S2A for mean values. No significant differences between the different molecules for all four states, using Mann-Whitney U tests.

**C.** Average proportion of DDMap populations per nucleus of CSL (green), Mam (red) and Hairless (brown) molecules in Notch-Off (-) or Notch-On (+) conditions as indicated. Nuclei as in Figure 2A. See Table S1B for mean values, Table S3C for p-values of two sample t-tests.

**Supplementary Figure S2: Recruitment and change in DDMap proportions of CSL complexes near *E(spl)-C* in Notch active conditions.**

**A.** Nuclear localisation of all vbSPT D1 molecule trajectories from the experiment in Figure 3A. Arrowhead shows location of *E(spl)-C* locus.

**B.** Nuclear localisation of all sub-diffusive molecule trajectories identified by DDMap analysis from the experiment in Figure 3A. Arrowhead shows location of *E(spl)-C* locus.

**C.** Average proportion of Brownian and sub-diffusive trajectories near *E(spl)-C* versus away for CSL (green), Mam (magenta) and Hairless (brown) in Notch-On conditions. Nuclei as in Figure 3A. Values plotted were calculated as (Proportion near- Proportion away)/Proportion away, hence a value greater than 0 represents an enrichment of the respective population near the target locus. Wilcoxon signed rank-sum tests were performed (* for $p<0.05$, ** for $p<0.01$, *** for $p<0.001$). See p-values Table S3F for p-values.

**Supplementary Figure S3: Clustering analysis – nearest neighbour histogram and anisotropic behaviour of CSL near bound clusters in Notch-Off conditions.**

**A.** Example of distance to $k^{th}$ nearest neighbour histogram for $k = 11$, for bound molecules (D1 and D2 populations) of a representative CSL Notch-On experiment. Bimodal distribution indicates that some molecules are in clusters (their $k^{th}$ neighbour is nearby) while others are not (their $k^{th}$ neighbour is further away).

**B.** Proportion of near and away diffusive jumps (D3 and D4) that were anisotropic for CSL in Notch-Off nuclei. Mean values: $0.238 \pm 0.013$ Control, $0.350 \pm 0.033$ Near, $0.222 \pm 0.013$ Away. Lines connect data from the same experiment (see Results & Methods for details on control analysis). Wilcoxon signed-rank tests were performed, $p = 0.016$ for Control vs Near and Near vs Away.

### Supplementary 1

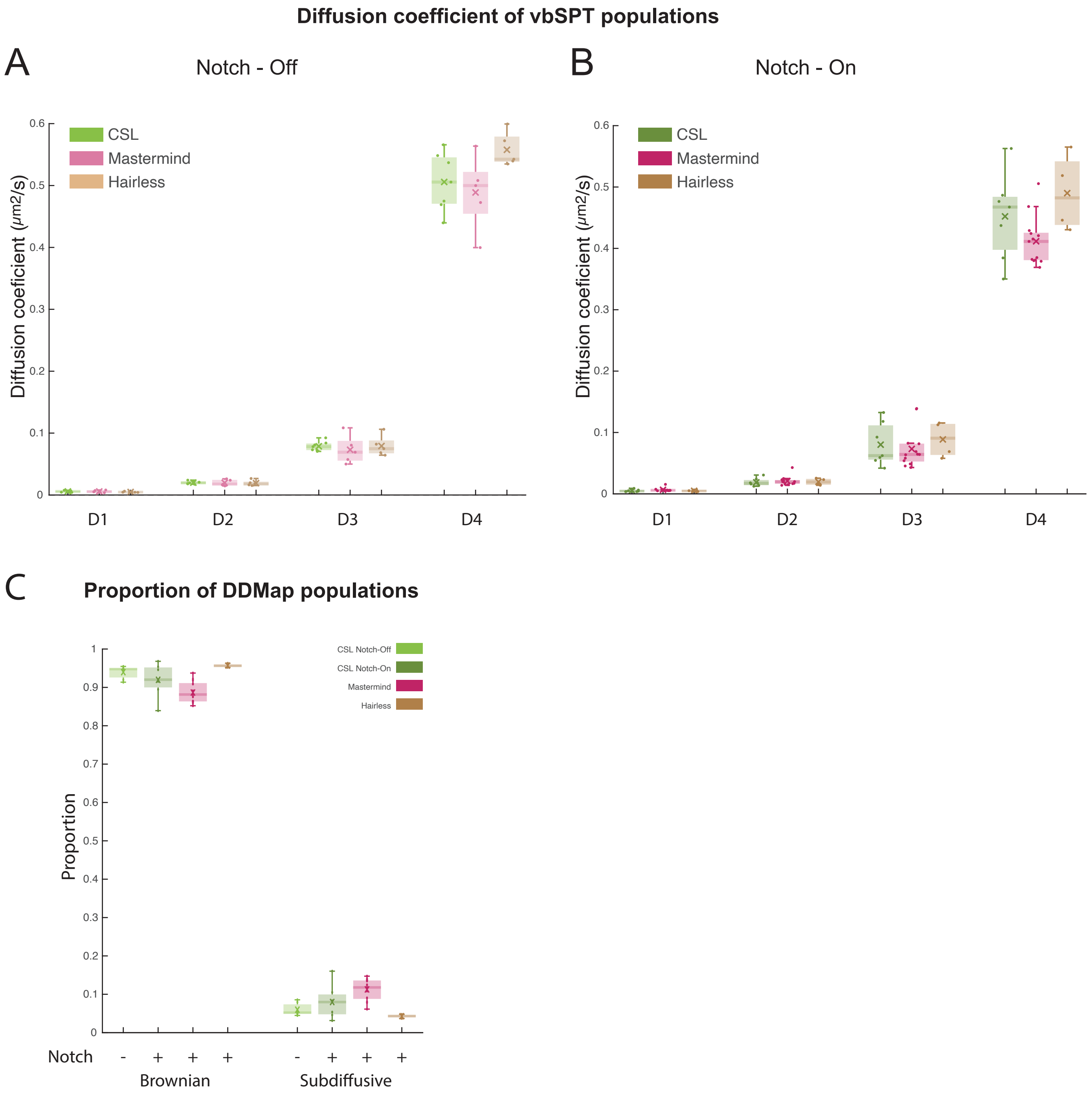

### Supplementary 2

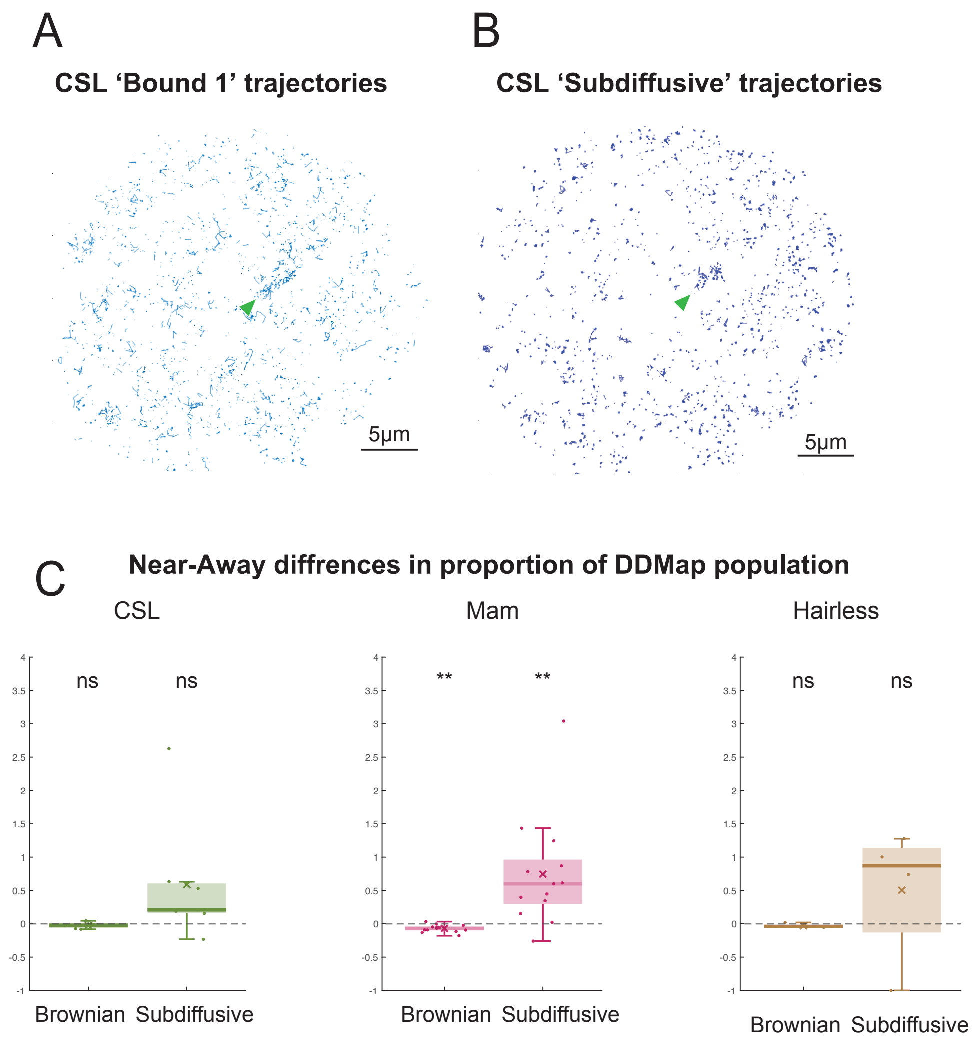

### Supplementary 3

A

**Bound CSL molecules' distance to 11th nearest neighbour**

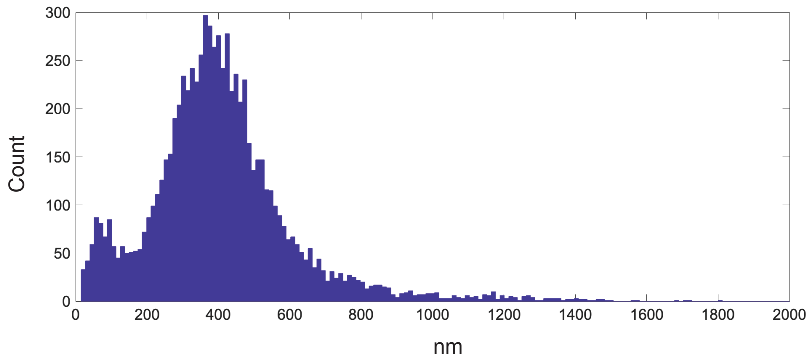


B

**Anisotropic behaviour of CSL near bound clusters**

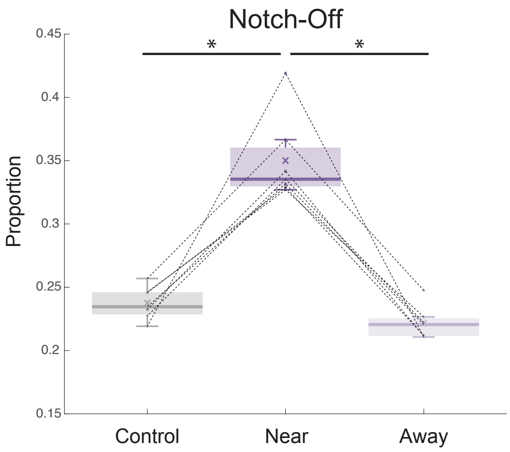

#### Supplementary Tables

**Table S1: Proportions of molecules assigned to each population.**

Table S1A: Mean values (± SD) of proportions of vbSPT populations (Fig 2A).

| | **CSL Notch-Off** | **CSL Notch-On** | **Mam Notch-On** | **Hairless Notch-On** |
|---|---|---|---|---|
| **D1** | 0.057 ± 0.026 | 0.090 ± 0.025 | 0.092 ± 0.053 | 0.083 ± 0.005 |
| **D2** | 0.187 ± 0.031 | 0.220 ± 0.034 | 0.345 ± 0.094 | 0.183 ± 0.062 |
| **D3** | 0.275 ± 0.095 | 0.370 ± 0.094 | 0.316 ± 0.129 | 0.203 ± 0.035 |
| **D4** | 0.481 ± 0.098 | 0.320 ± 0.116 | 0.247 ± 0.073 | 0.531 ± 0.082 |

Table S1B: Mean values (± SD) of proportions of DDMap populations (Fig Supp 1C).

| | **CSL Notch-Off** | **CSL Notch-On** | **Mam Notch-On** | **Hairless Notch-On** |
|---|---|---|---|---|
| **Brownian** | 0.940 ± 0.016 | 0.920± 0.043 | 0.887 ± 0.028 | 0.957 ± 0.005 |
| **Sub-diffusion** | 0.060 ± 0.016 | 0.080 ± 0.043 | 0.113 ± 0.028 | 0.043 ± 0.005 |

**Table S2: Diffusion constants.**

Table S2A: Mean values (± SD) of diffusion coefficients of vbSPT populations (Fig S1A, S1B).

| **Notch-Off** | **CSL** | **Mam** | **Hairless** |
|---|---|---|---|
| **D1** | 0.006 ± 0.001 | 0.006 ± 0.002 | 0.005 ± 0.001 |
| **D2** | 0.020 ± 0.002 | 0.020 ± 0.005 | 0.019 ± 0.005 |
| **D3** | 0.079 ± 0.008 | 0.073 ± 0.023 | 0.079 ± 0.017 |
| **D4** | 0.506 ± 0.047 | 0.489 ± 0.060 | 0.558 ± 0.028 |
| **Notch-On** | **CSL** | **Mam** | **Hairless** |
| **D1** | 0.005 ± 0.002 | 0.007 ± 0.003 | 0.005 ± 0.002 |
| **D2** | 0.019 ± 0.006 | 0.021 ± 0.007 | 0.020 ± 0.006 |
| **D3** | 0.080 ± 0.035 | 0.073 ± 0.032 | 0.089 ± 0.030 |
| **D4** | 0.452 ± 0.070 | 0.412 ± 0.040 | 0.490 ± 0.063 |

Table S2B: Mean values (± SD) of diffusion coefficients of DDMap populations (Fig 2B).

| | **CSL Notch-Off** | **CSL Notch-On** | **Mam Notch-On** | **Hairless Notch-On** |
|---|---|---|---|---|
| **Brownian** | 0.317 ± 0.066 | 0.219 ± 0.076 | 0.161 ± 0.030 | 0.314 ± 0.073 |
| **Sub-diffusion** | 0.036 ± 0.007 | 0.026 ± 0.005 | 0.027 ± 0.006 | 0.024 ± 0.005 |

**Table S3: Results from statistical tests.**

Table S3A: p-values for comparisons of proportions of vbSPT populations (Fig 2A).

| CSL Notch-Off | CSL Notch-On | Mann-Whitney U test p-value |
|---|---|---|
| D1 | D1 | 0.026 |
| D2 | D2 | 0.097 |
| D3 | D3 | 0.128 |
| D4 | D4 | 0.018 |
| **CSL Notch-On** | **Mam Notch-On** | **Mann-Whitney U test p-value** |
| D1 | D1 | 0.939 |
| D2 | D2 | 0.002 |
| D3 | D3 | 0.211 |
| D4 | D4 | 0.157 |
| **CSL Notch-On** | **Hairless Notch-On** | **Mann-Whitney U test p-value** |
| D1 | D1 | 0.315 |
| D2 | D2 | 0.412 |
| D3 | D3 | 0.006 |
| D4 | D4 | 0.024 |

Table S3B : p-values for comparisons of diffusion coefficients of DDMap populations (Fig 2B).

| CSL Notch-Off | CSL Notch-On | Two-sample t-test p-value |
|---|---|---|
| Brownian | Brownian | 0.023 |
| Sub-diffusion | Sub-diffusion | 0.0134 |
| **CSL Notch-On** | **Mam Notch-On** | **Two-sample t-test p-value** |
| Brownian | Brownian | 0.103 |
| Sub-diffusion | Sub-diffusion | 0.706 |
| **CSL Notch-On** | **Hairless Notch-On** | **Two-sample t-test p-value** |
| Brownian | Brownian | 0.068 |
| Sub-diffusion | Sub-diffusion | 0.489 |

Table S3C: p-values for comparisons of proportions of DDMap populations (Fig S1C).

| **CSL Notch-Off** | **CSL Notch-On** | **Two-sample t-test p-value** |
|---|---|---|
| Brownian | Brownian | 0.286 |
| Sub-diffusion | Sub-diffusion | 0.285 |
| **CSL Notch-On** | **Mam Notch-On** | **Two-sample t-test p-value** |
| Brownian | Brownian | 0.050 |
| Sub-diffusion | Sub-diffusion | 0.052 |
| **CSL Notch-On** | **Hairless Notch-On** | **Two-sample t-test p-value** |
| Brownian | Brownian | 0.065 |
| Sub-diffusion | Sub-diffusion | 0.065 |

Table S3D: p-values for Near and Away comparison of proportions of vbSPT populations. Statistical significance indicates difference from 0 of Near-away ratio (Fig 3E).

| **CSL Notch-On** | **One-sample t-test p-value** |
|---|---|
| D1 | 0.004 |
| D2 | 0.139 |
| D3 | 0.238 |
| D4 | 0.003 |
| **Mam Notch-On** | **One-sample t-test p-value** |
| D1 | 0.022 |
| D2 | 0.119 |
| D3 | 0.199 |
| D4 | 1.06E-05 |
| **Hairless Notch-On** | **One-sample t-test p-value** |
| D1 | 0.301 |
| D2 | 0.032 |
| D3 | 0.926 |
| D4 | 0.003 |

Table S3E: p-values for Near andAway comparison  of diffusion coefficients of DDMap populations. Statistical significance indicates difference from 0 of Near-away ratio (Fig 3F).

| **CSL Notch-On** | **Wilcoxon signed rank-sum test p-value** |
|---|---|
| Brownian | 0.016 |
| Sub-diffusion | 0.813 |
| **Mam Notch-On** | **Wilcoxon signed rank-sum test p-value** |
| Brownian | 0.003 |
| Sub-diffusion | 0.685 |
| **Hairless Notch-On** | **Wilcoxon signed rank-sum test p-value** |
| Brownian | 0.125 |
| Sub-diffusion | 1.000 |

Table S3F: p-values for Near and Away comparisons of proportions of DDMap populations. Statistical significance indicates difference from 0 of Near-away ratio (Fig S2C).

| **CSL Notch-On** | **Wilcoxon signed rank-sum test p-value** |
|---|---|
| Brownian | 0.156 |
| Sub-diffusion | 0.109 |
| **Mam Notch-On** | **Wilcoxon signed rank-sum test p-value** |
| Brownian | 0.001 |
| Sub-diffusion | 0.001 |
| **Hairless Notch-On** | **Wilcoxon signed rank-sum test p-value** |
| Brownian | 0.250 |
| Sub-diffusion | 0.375 |

**Table S4: Genotypes of flies used for each condition.**

| | |
|---|---|
| H2-AV | H2-AV-Halo |
| CSL Notch-Off | 1151-Gal4;; Su(H)-Halo X UAS-LacZ ; E(spl)mdelta[IntB], UAS-P31B::GFP |
| CSL Notch-On | 1151-Gal4;; Su(H)-Halo X UAS-NΔECD ; E(spl)mdelta[IntB], UAS-P31B::GFP |
| Mam Notch-Off | 1151-Gal4; Mam-Halo X UAS-LacZ ; E(spl)mdelta[IntB], UAS-P31B::GFP |
| Mam Notch-On | 1151-Gal4; Mam-Halo X UAS-NΔECD ; E(spl)mdelta[IntB], UAS-P31B::GFP |
| Hairless Notch-Off | 1151-Gal4; Hairless-Halo X UAS-LacZ ; E(spl)mdelta[IntB], UAS-P31B::GFP |
| Hairless Notch-On | 1151-Gal4; Hairless-Halo X UAS-NΔECD ; E(spl)mdelta[IntB], UAS-P31B::GFP |
